# Movements of ancient human endogenous retroviruses detected in SOX2-expressing cells

**DOI:** 10.1101/2020.07.14.202135

**Authors:** Kazuaki Monde, Yorifumi Satou, Mizuki Goto, Yoshikazu Uchiyama, Jumpei Ito, Taku Kaitsuka, Hiromi Terasawa, Shinya Yamaga, Tomoya Matsusako, Fan-Yan Wei, Ituro Inoue, Kazuhito Tomizawa, Akira Ono, Takumi Era, Tomohiro Sawa, Yosuke Maeda

## Abstract

Human endogenous retroviruses (HERVs) occupy approximately 8% of human genome. HERVs, which are transcribed in early embryos, are epigenetically silenced in somatic cells, except in pathological contexts. HERV-K is thought to protect the embryo from exogenous viral infection. However, uncontrollable HERV-K expression in somatic cells has been implicated in several diseases. Here, we show that SOX2, which plays a key role in maintaining pluripotency of stem cells, is critical for the transcription of HERV-K LTR5Hs. HERV-K can undergo retrotransposition within producer cells in the absence of Env expression. Furthermore, new HERV-K integration sites were identified in a long-term culture of induced pluripotent stem cells, which express SOX2. Together, these results suggest the possibility that the strict dependence of HERV-K on SOX2 have allowed contribution of HERV-K to the protection of early embryos during evolution while limiting potentially harmful effects of HERV-K retrotransposition on host genome integrity to these early embryos.

## Introduction

Endogenous retroelements are mobile genetic elements that constitute more than 40% of the human genome. Human endogenous retroviruses (HERVs), which encode the long terminal repeat (LTR)-containing elements, occupy about 8% of the human genome (Bannert & Kurth, 2004; Lander et al, 2001; Venter et al, 2001). For more than 20 million years, HERVs that have persisted in germ-cell lineages have been transmitted vertically from ancestors to descendant (Boeke & Stoye, 1997). At present, almost all HERVs have acquired numerous mutations or deletions. However, HERV-K, a relatively new endogenous retrovirus, apparently encodes intact open reading frames in the human genome (Turner et al, 2001), although no replication-competent HERV-K has been detected (Beimforde et al, 2008; Boller et al, 2008; Lee & Bieniasz, 2007; Stoye, 2012). HERV-K is transcribed during early embryogenesis or exogenous viral infection, producing HERV-K proteins that appear to protect the host cells from viral attack (Grow et al, 2015; Monde et al, 2012; Monde et al, 2017; Terry et al, 2017). HERV-K expression has also been noted in various human diseases, including autoimmune disorders, neurological diseases, infectious diseases, and cancer (Young et al, 2013).

Long interspersed nuclear elements (LINE-1), which are classified among the non-LTR retroelements, are transposition competent (Beck et al, 2010; Brouha et al, 2003; Mills et al, 2007). The transposition of LINE-1 mainly occurs in germ cells during early embryonic development. These transposition events might cause pathogenesis by altering the structures, expression, and functions of genes (Beck et al, 2011; Han et al, 2004; Hancks & Kazazian, 2012). Therefore, transposition is regulated by histone modifications and DNA methylation to avoid the harmful mutations in the genomes (Bourc’his & Bestor, 2004; Levin & Moran, 2011). Recent advances in sequencing technology have allowed the detection of non-reference HERV-K, which is absent from the human genome sequence, in the population (Wildschutte et al, 2016), although HERV-K retrotransposition activity has not yet been reported.

HERV-K encodes the 5’-LTR and 3’-LTR at the upstream and downstream of the viral protein ORFs respectively. HERV-K LTR has preserved their promoter activity, and HERV-K is transcribed in embryonic stem cells, several cancer cells, and virus-infected T cells (Grow et al, 2015). The transcription factors Sp1 and Sp3 drive HERV-K transcription in teratocarcinoma cells and melanoma cells (Fuchs et al, 2011). The melanoma-specific transcription factor MITF-M is also required for the activation of the HERV-K LTR (Katoh et al, 2011). In virus-infected cells, viral transcription factors Tat and Tax are associated with HERV-K expression (Gonzalez-Hernandez et al, 2012; Toufaily et al, 2011). In embryonic stem cells, DNA hypomethylation and OCT3/4-binding to the HERV-K LTR synergistically facilitate HERV-K transcription (Grow et al, 2015). However, it remains unclear whether these transcription factors are essential for HERV-K activation.

Here, we show that SOX2, rather than OCT3/4, is the major factor for activating the transcription of HERV-K LTR5Hs, which is the youngest HERV-K subfamily (Turner et al, 2001). Consistent with this finding, a large amount of HERV-K Gag is expressed in induced pluripotent stem (iPS) cells, which are SOX2-expressing cells. We used next-generation sequencing (NGS) to analyze the genomes of iPS cells and determined the HERV-K integration sites. Surprisingly, we found that new HERV-K insertions into the genome increase in a manner dependent upon the culture period, suggesting that HERV-K retrotransposition occurs in SOX2-expressing cells. Our results suggest that HERV-K is not a harmless fossil left in the human genome; rather, it retains the ability to spread among the human genome by retrotransposition, but is normally repressed due to its dependence on SOX2 expression.

## Results

### SOX2 activates HERV-K transcription

Teratocarcinomas are germ-cell tumors, and teratocarcinoma cells constitutively express HERV-K proteins and release HERV-K particles from their plasma membranes (Bieda et al, 2001; Boller et al, 1983). Several transcription factors, including MITF, MZF1, NF-Y, GATA-2, and OCT3/4, are required to activate the HERV-K LTR (Grow et al, 2015; Katoh et al, 2011; Yu et al, 2005). Wysocka’s research group identified the consensus OCT3/4-binding motifs in HERV-K LTR5Hs, and demonstrated the transcriptional activation of HERV-K by OCT3/4 in human preimplantation embryos (Grow et al, 2015). However, it is unknown whether the expression of OCT3/4 is sufficient for the transcriptional activation of HERV-K. Here, we identified the region of HERV-K responsible for the transcription of HERV-K LTR5Hs using deletion mutants of HERV-K LTR5Hs in teratocarcinoma cells (NCCIT cells) (Supplemental Fig. S1A). These results show that the deletion of nucleotides nt 650–700 in LTR5Hs causes the loss of its transactivation activity (Supplemental Fig. S1B and S1C). With the PROMO software (Farre et al, 2003; Messeguer et al, 2002), which is used to identify putative transcription factors, we identified 15 SOX2-binding motifs (#1–#12) and two OCT3/4-binding motifs (#13 and #14) in LTR5Hs (Fig. 1A and Supplemental Fig. S1). Some SOX2-binding motifs overlapped with each other (#10 and #11), and therefore these motifs were called each same number. Two OCT3/4-binding motifs (#13 and #14) and three SOX2-binding motifs (#9, #10, and #11) occurred in the region nt 650–700 in LTR5Hs (Fig. 1A and Supplemental Fig. S1A). Based on a chromatin immunoprecipitation (ChIP) analysis database in embryonic stem cells, there were two peaks of SOX2 binding at nt 200 and 700 of the HERV-K LTR genome (Fig. 1B). OCT3/4-binding peaks were similar to those for SOX2 in the HERV-K LTR genome. To determine whether OCT3/4 is sufficient for the transactivation of HERV-K LTR, as reported previously (Grow et al, 2015), we cotransfected HeLa cells with plasmids encoding each transcription factor (OCT3/4, SOX2, KLF4, NANOG) and the HERV-K LTR-Luc (Fig. 1C). Unexpectedly, we found that OCT3/4 was not sufficient to activate the transcription of HERV-K LTR in HeLa cells. The transcription of LTR mutants, with mutations in the OCT3/4-binding motifs, was slightly reduced, but not significantly so, in NCCIT cells (Supplemental Fig. S1D) and when SOX2, KLF4 and OCT3/4 were overexpressed in HeLa cells (Supplemental Fig. S1E). In contrast, SOX2 markedly activated HERV-K LTR transcription (Fig. 1C). In the presence of SOX2, KLF4 slightly increased the transactivation of HERV-K, but OCT3/4 reduced the effect of SOX2 (Supplemental Fig. S1F). In the presence of both SOX2 and KLF4, OCT3/4 increased the transactivation of HERV-K. The transactivation of HERV-K was dose-dependently enhanced by expression of SOX2 alone (Fig. 1D and 1E). However, OCT3/4 alone did not alter the transactivation of HERV-K, even when overexpressed.

**Fig. 1.**
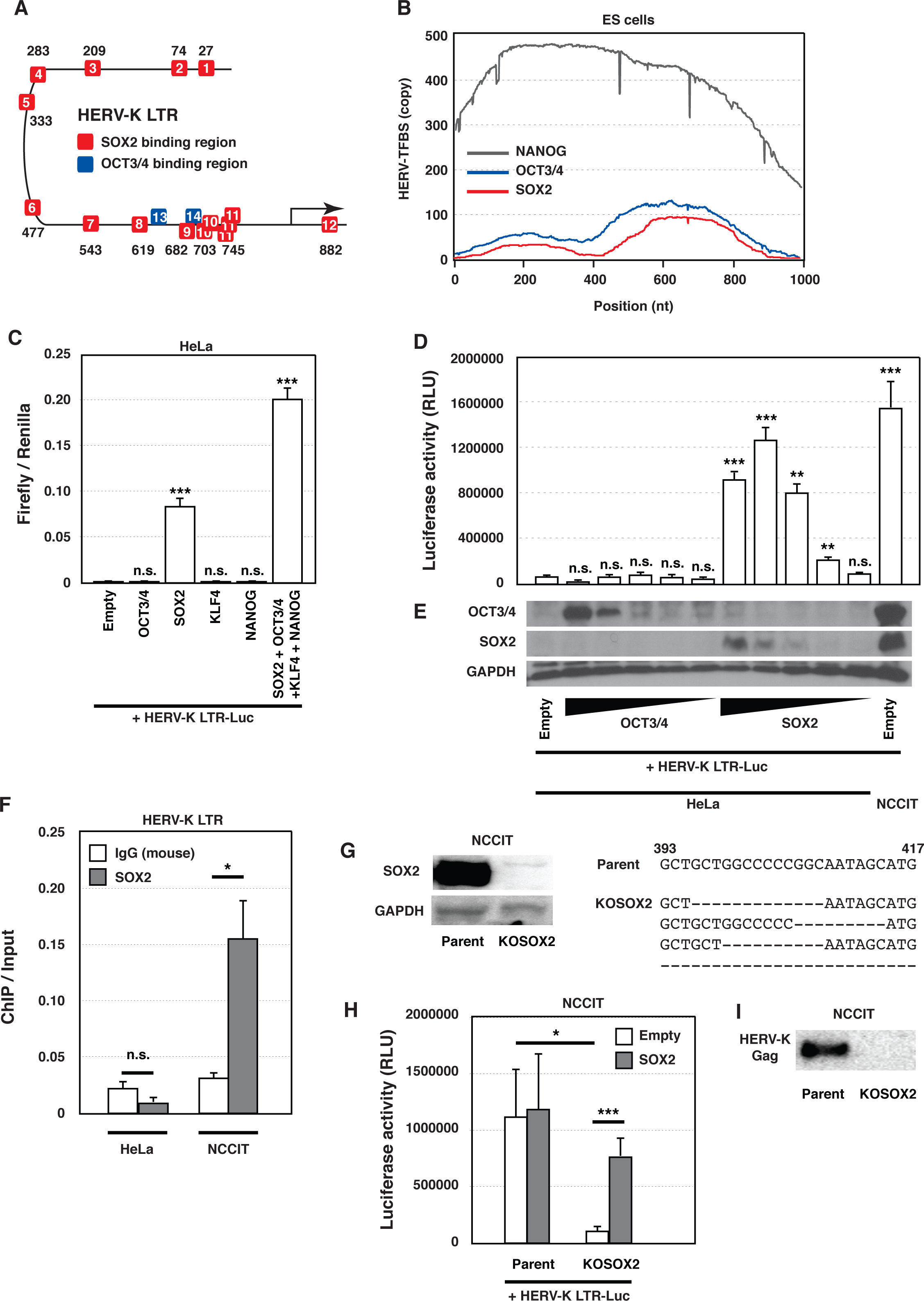
SOX2 contributes to the promoter function of HERV-K LTR. (A) SOX2 and OCT3/4 binding motifs were identified in the HERV-K LTR with the PROMO software, which is used to identify transcription factor binding motifs. (B) Binding sites of SOX2, OCT3/4 and NANOG on respective LTR5Hs copies in human ES cells were collected from ENCODE ChIP-Seq dataset, and the positions on the consensus sequence of LTR5Hs are shown. (C) HeLa cells were cotransfected with the plasmid pHERV-K_CON_ LTR-Luc, the indicated plasmids, and the *Renilla*-Luc plasmid. The firefly and renilla luciferase activities were measured. (D) HeLa cells were cotransfected with pHERV-K LTR-Luc and different amounts of the indicated plasmids. The luciferase activity was measured. (E) Amounts of OCT3/4, SOX2, and GAPDH proteins were measured with western blotting. (F) Chromatins in HeLa and NCCIT cells were extracted, and SOX2-binding DNA fragments were immunoprecipitated with the indicated antibodies. HERV-K LTRs in the immunoprecipitated DNA were measured with qPCR. (G) Amounts of SOX2 and GAPDH proteins in NCCIT and SOX2-knockout NCCIT (NCCIT/KOSOX2) cells were measured with western blotting. The sequences of SOX2 in each cell were analyzed. (H) NCCIT and NCCIT/KOSOX2 cells were cotransfected with pHERV-K LTR-Luc and SOX2-encoding plasmids. The luciferase activity was measured. (C, D, F, and H) Data from three independent experiments are shown as means ± standard deviations. *P* values were determined with Student’s *t* test. \**P* < 0.01; \*\**P* < 0.001; \*\*\**P* < 0.0001; n.s., not significant. (I) Amounts of mature HERV-K Gag in the supernatants of NCCIT and NCCIT/KOSOX2 cells were measured with western blotting.

Because NCCIT cells express large amounts of endogenous SOX2 (Fig. 1E), we examined the binding of endogenous SOX2 to chromosomal HERV-K LTR with a ChIP assay (Fig. 1F). The results showed that endogenous SOX2 binds to the chromosomal HERV-K LTR in NCCIT cells. To confirm that endogenous SOX2 drives HERV-K transcription, we established SOX2-knockout NCCIT cells (NCCIT/KOSOX2) (Fig. 1G). Although the genome of the NCCIT/KOSOX2 cells encodes four different sequence patterns, no intact *SOX2* gene was detected in the NCCIT/KOSOX2 cells (Fig. 1G right). HERV-K LTR transactivation was dramatically reduced in the NCCIT/KOSOX2 cells, but not completely lost, and was rescued by the transfection of SOX2 (Fig. 1H). The mature HERV-K Gag protein (37 kDa) in the viral particles disappeared from the supernatant of the KOSOX2 cells (Fig. 1I).

Together, these results indicate that SOX2 is an essential transcription factor for expression of HERV-K LTR5Hs, and that both OCT3/4 and KLF4 drive HERV-K transcription in the presence of SOX2.

### Multiple SOX2-binding motifs activate the HERV-K transcription

With the Promo software and the ChIP database, we localized nine of 14 SOX2-binding motifs (#3, #4, #7, #8, #9, #10, and #11) around nt 200 and 700 of the HERV-K LTR genome (Fig. 1A and 1B). Based on Fig. S1C, we speculated that a deletion of the single SOX2-binding motif #9 might abolish the transactivation of HERV-K LTR. To determine the region responsible for HERV-K transactivation by SOX2, HeLa cells were cotransfected with plasmids encoding HERV-K LTR–luc mutants (del#01-#12) and SOX2, KLF4, and OCT3/4. Unexpectedly, any single deletion of a SOX2-binding motif did not reduce the transactivation of HERV-K LTR in HeLa cells (Fig. 2A) or NCCIT cells (Fig. 2B). However, the deletion of all SOX2-binding motifs dramatically reduced HERV-K transactivation in both HeLa and NCCIT cells. Notably, LTR sequences that contain some single deletions showed similar activity to that of WT HERV-K LTR, but other single deletions enhanced the LTR activity, suggesting the redundancy and/or interference between SOX2-binding motifs. Therefore, we designed mutants of LTR5Hs with multiple deletions of SOX2-binding motifs (Fig. 2C and 2D). Deletion of SOX2-binding motifs #03, #08, #09, and #10 around nt 200 and 700 in LTR5Hs, which correspond to two major SOX2-binding regions (Fig. 1A and 1B), reduced HERV-K transactivation to the same degree as the deletion of all the SOX2-binding motifs in both HeLa cells (Fig. 2C) and NCCIT cells (Fig. 2D). These results suggest that SOX2 activates HERV-K transcription even after the accumulation of several mutations in LTR5Hs during its biological evolution.

**Fig. 2.**
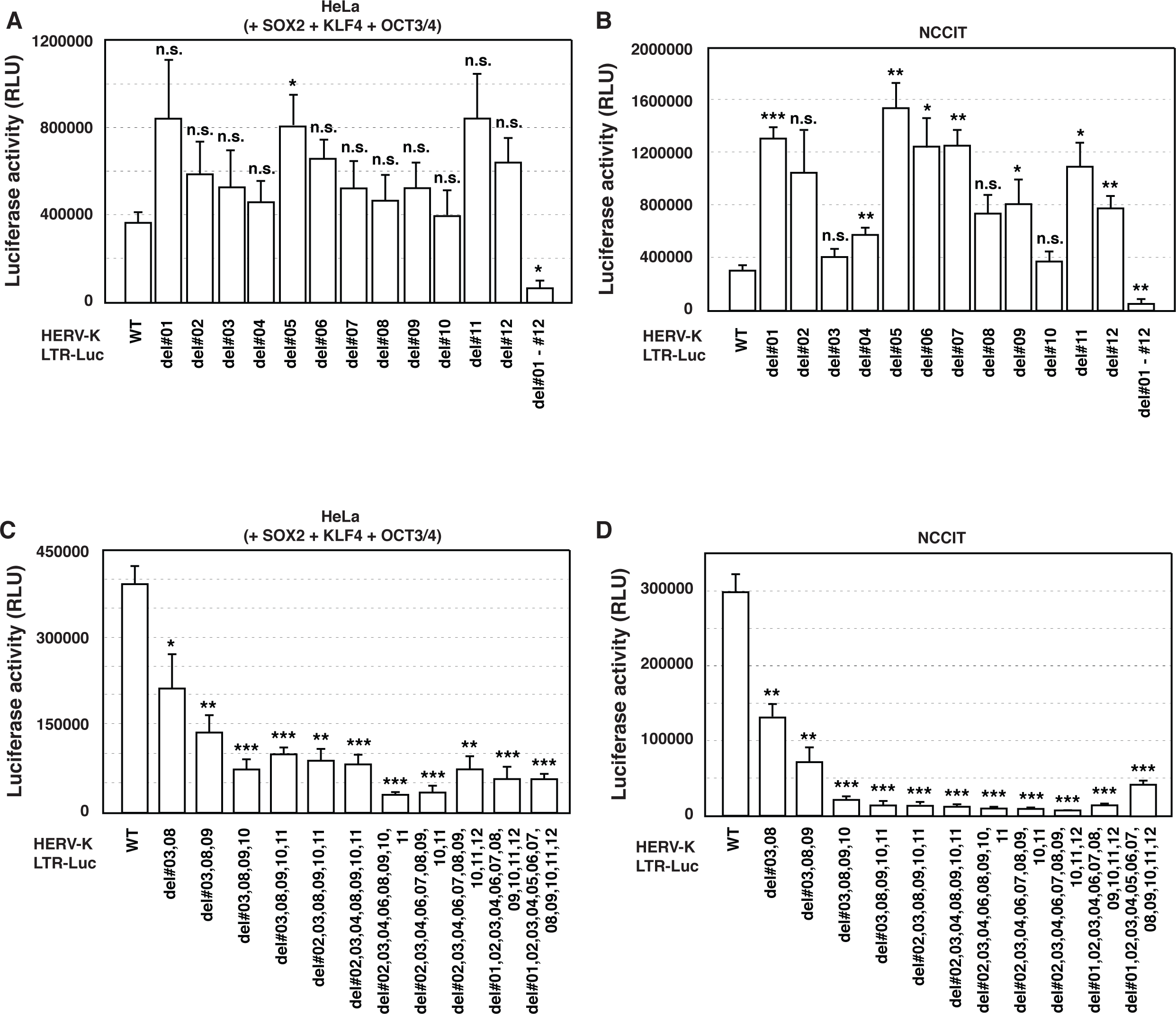
Multiple SOX2-binding motifs contribute to HERV-K transcription. HeLa (A and C) and NCCIT cells (B and D) were cotransfected with pHERV-K LTR mutants and the indicated plasmids. The luciferase activity was measured. (A, B, C, and D) Data from three independent experiments are shown as means ± standard deviations. *P* values were determined with Student’s *t* test. \**P* < 0.01; \*\**P* < 0.001; \*\*\**P* < 0.0001; n.s., not significant.

### SOX2 activates chromosomal HERV-K expression

HERV-K genomes have a CpG island between the LTR and the *gag* gene (Fig. 3A), which is hypermethylated in HeLa cells (Fig. 3C) compared with NCCIT cells (Fig. 3B). This suggests that HERV-K genomes are packed into heterochromatin and are silenced in HeLa cells. To confirm the modification of the chromatin, we treated HeLa cells with 5-aza-2’-deoxycytidine to hypomethylate the genome. The hypomethylation of the genome enhanced HERV-K Gag mRNA expression when SOX2 was overexpressed in the HeLa cells (Fig. 3D). These results indicate that DNA hypomethylation and SOX2 expression synergistically induce the expression of HERV-K genes in the human genome.

**Fig. 3.**
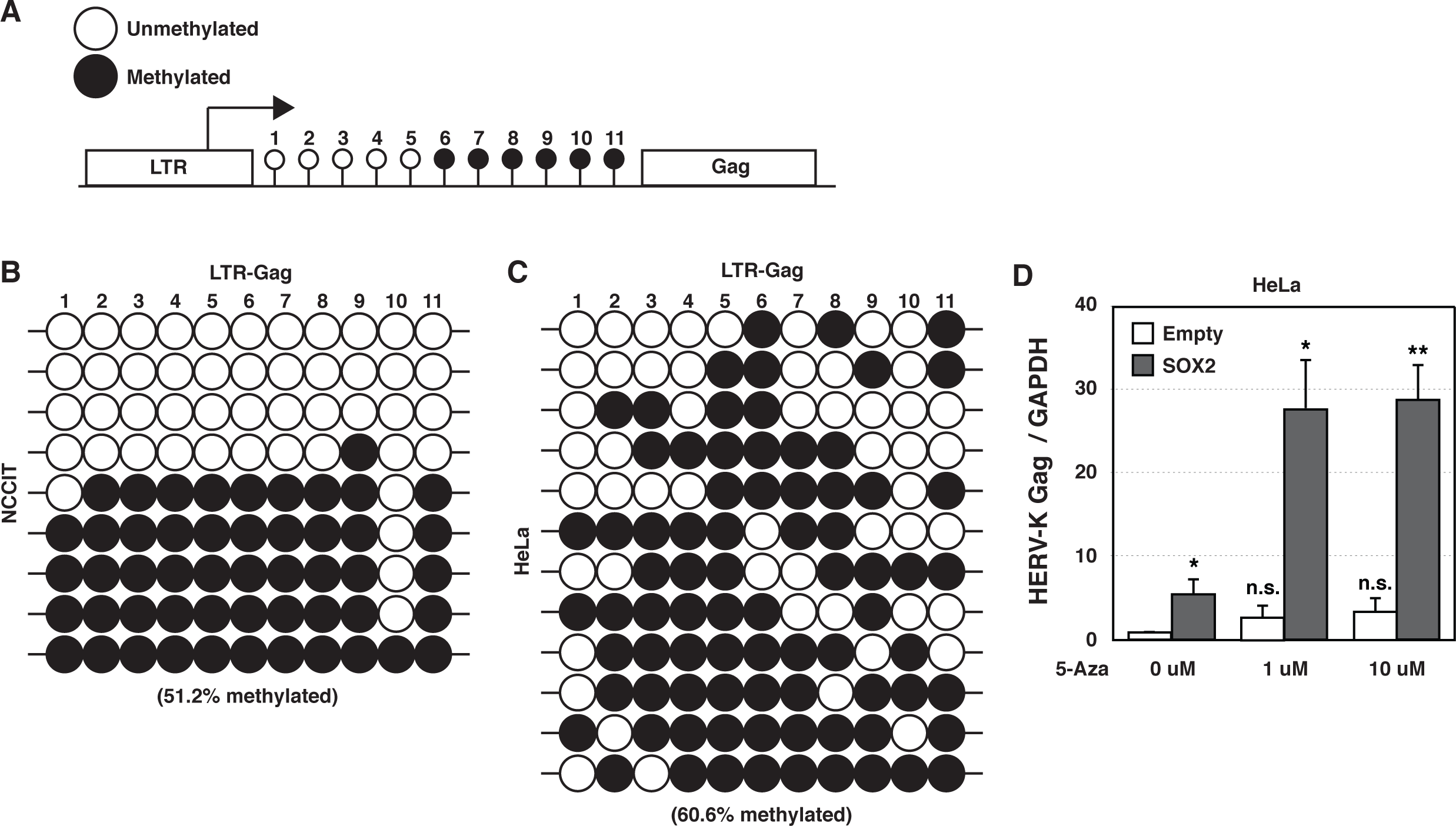
HERV-K genome is hypermethylated in HeLa cells. (A) There is likely to be a CpG island (11 CG nucleotides) between HERV-K LTR and *gag*. (B and C) DNA was extracted from NCCIT (B) and HeLa cells (C). The sequences of nine HERV-K genomes between LTR and Gag in NCCIT cells and 12 HERV-K genomes in HeLa cells were analyzed after the DNAs were treated with bisulfite to convert cytosine residues to uracil. White circles indicate unmethylated nucleotides and black circles indicate methylated nucleotides in the CpG island. (D) HeLa cells were treated with 5-aza-2’-deoxycytidine for 1 day, and then transfected with a plasmid encoding SOX2. Two days after transfection, the amounts of HERV-K Gag mRNA were measured with RT–qPCR. Data from three independent experiments are shown as means ± standard deviations. *P* values were determined using Student’s *t* test. \**P* < 0.01; \*\**P* < 0.001; n.s., not significant.

### SOX2 activates the 5’ and 3’ LTR5Hs of HERV-K

Because LTR sequences of HERV-K is classified into three major groups (LTR5Hs, 5A, and 5B), we cloned 18 different HERV-K LTRs from NCCIT cells and investigated whether HERV-K LTR transactivation by SOX2 is conserved among the three different groups. The LTR sequences of the HERV-K 5Hs group (LTR5Hs) are the part of the most recently integrated sequences (around 9.1 million years ago) (Subramanian et al, 2011). There are two types of LTR5Hs proviruses that are classified based on the presence (type 1) or absence (type 2) of a 292 bp deletion at the pol-env junction. The LTRs of 5A and 5B groups (LTR5A and LTR5B) are associated with proviruses that are mainly classified as type 2 (Subramanian et al, 2011). The LTR5B proviruses include the oldest insertions (around 27.9 million years ago), and LTR5A proviruses (around 20.1 million years ago) are originating from LTR5B at an estimated-standard mutation rate of 0.24-0.45% per million years based on the LTR-based and internal-based phylogenies (Subramanian et al, 2011). Interestingly, both 5’- and 3’-LTR of LTR5Hs and three out of four LTR5B were significantly activated by SOX2, whereas three out of four LTR5A were no activated in SOX2-expressing HeLa cells (Fig. 4A) and NCCIT cells (Fig. 4B). A phylogenetic analysis of HERV-K LTRs showed that SOX2-responsive HERV-K LTRs are closely related (Fig. 4C). Both the newest and oldest HERV-K LTRs integrated into genomes retain the capacity for SOX2-dependent transactivation, suggesting that acquiring or maintaining this capacity is advantageous for coexistence between HERV-K and the host.

**Fig. 4.**
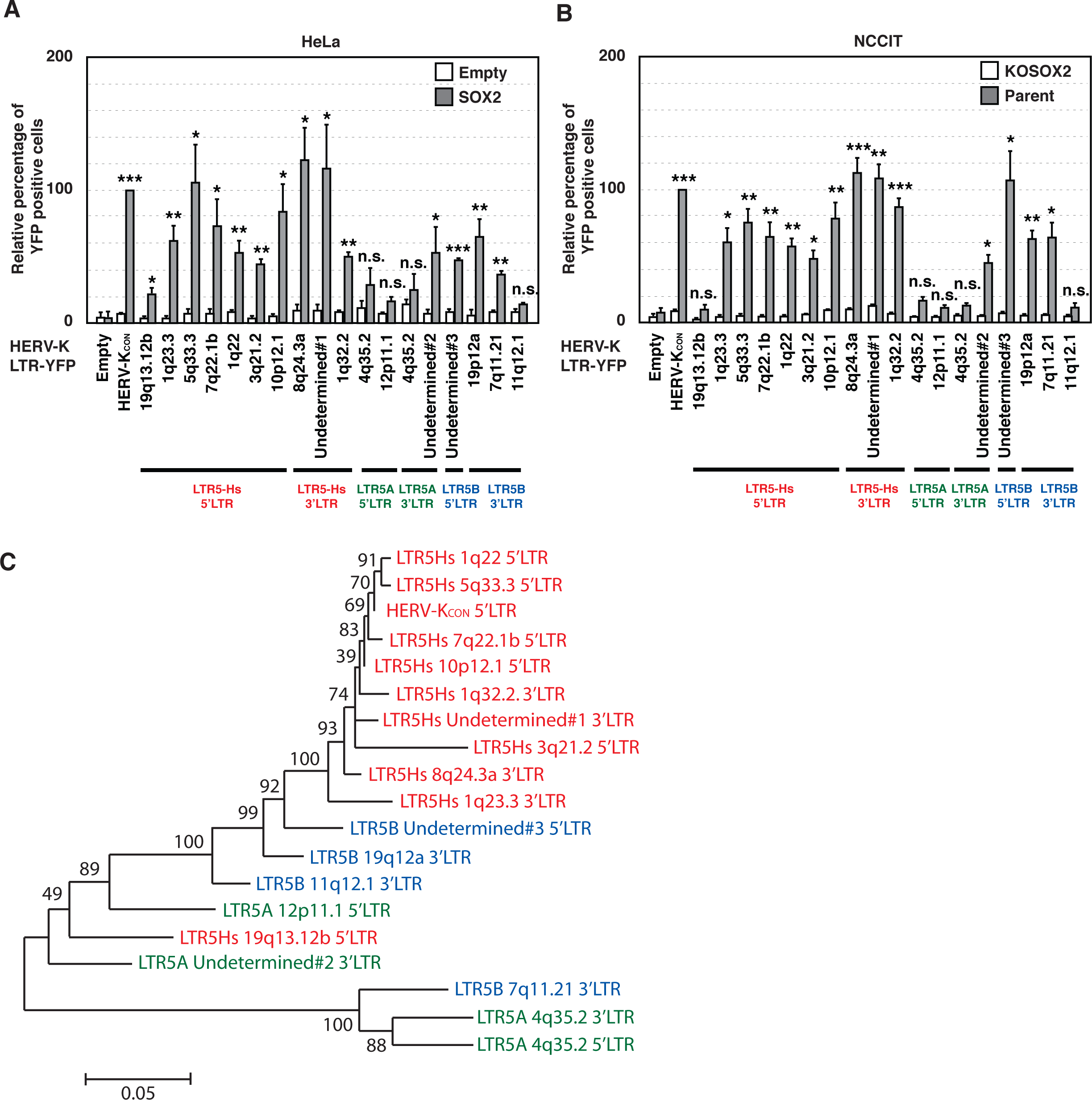
HERV-K transactivation by SOX2 is conserved among HERV-K LTR5Hs. HERV-K LTR series were amplified with PCR from genome into NCCIT cells and inserted upstream from the *YFP* gene. HeLa cells (A) were cotransfected with plasmid encoding SOX2 and the pHERV-K LTR–YFP series. NCCIT (Parent) and NCCIT/KOSOX2 cells (B) were transfected with the pHERV-K LTR-YFP series. The yellow fluorescent protein (YFP)-positive cells were analyzed with flow cytometry. (C) Neighbor-joining tree was constructed based on the aligned nucleotide sequences corresponding to HERV-K LTRs within NCCIT cells.

### Reconstructed HERV-K has retrotransposition activity

HERV-K LTR5Hs is expressed in SOX2-expressing cells, such as germ cells, but it is unclear whether HERV-K has retrotransposition activity within these cells. To examine the retrotransposition activity of HERV-K, we designed a HERV-K_CON_ construct encoding intron-inserting nanoluciferase (inNanoluc) (Fig. 5A). After the transcription of HERV-K from the *Cytomegalovirus* (CMV) promoter, the orientation of the intact reporter gene was reversed by splicing, and the CMV promoter at the 5’-UTR was then replaced with U3 through reverse transcription (Fig. 5A bottom). The reverse-transcribed HERV-K integrated into genome, and the intact reporter gene was transcribed from the *Simian virus 40* (SV40) promoter. The nanoluciferase values indirectly reflected the retrotransposition activity of HERV-K. HERV-K GagProPol, which encodes full-length *gag, pro*, and *pol*, showed nanoluciferase activity 5 days after transfection, whereas HERV-K deltaGagProPol, which encodes only truncated-*gag*, showed only slight nanoluciferase activity in HeLa cells (Fig. 5B). This suggests that HERV-K protease, reverse transcriptase, and/or integrase is required for HERV-K retrotransposition. Plasmids encoding HERV-K deltaProPol, a protease mutant (D203N), a reverse transcriptase mutant (SIAA), or an integrase mutant (DR1, DR2) with inNanoluc reporter gene were cotransfected into HeLa cells with or without a protein expression plasmid encoding HERV-K GagProPol (Fig. 5C). Although these mutants also showed faint nanoluciferase activity, expression of HERV-K GagProPol rescued the nanoluciferase activity of these mutants. This indicates that HERV-K protease, reverse transcriptase, and integrase are required for the retrotransposition of HERV-K. It also suggests that the assembly of intact proteases, reverse transcriptases, and integrases of different HERV-K origins can complement defective HERV-Ks during HERV-K retrotransposition.

**Fig. 5.**
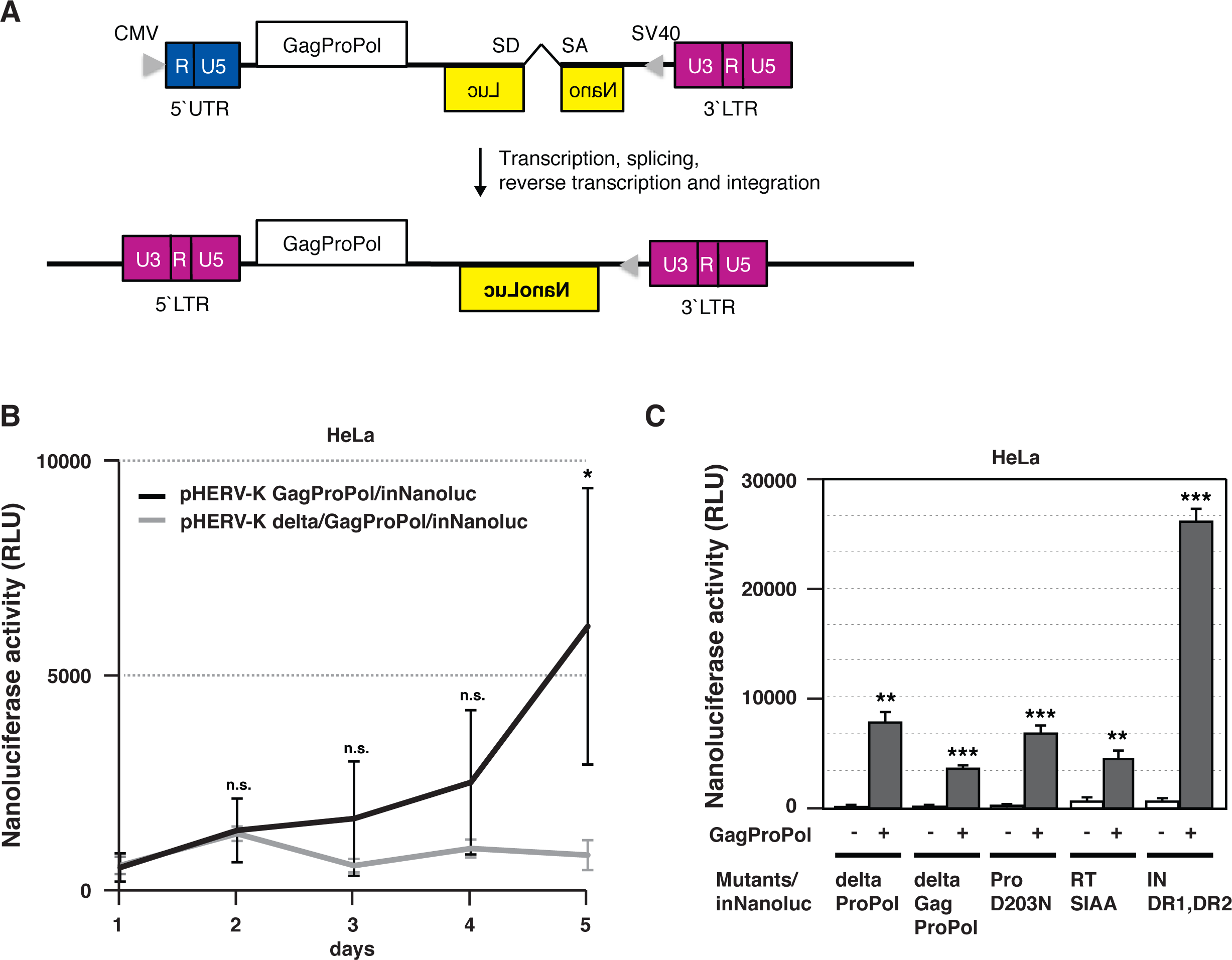
HERV-K has retrotransposition activity in HeLa cells. (A) The construction of pHERV-K GagProPol/inNanoluc is described. The 5’ U3 region was replaced with the CMV promoter. Intron-disrupted Nanoluc (inNluc) and SV40 were introduced into the Env region in an antisense orientation. (B) HeLa cells were transfected with pHERV-K GagProPol/inNanoluc or pHERV-K del/GagProPol/inNanoluc. Nanoluciferase activity was measured each day with a nanoluciferase reporter assay. (C) HeLa cells were transfected with pHERV-K mutants/inNanoluc alone or cotransfected with pHERV-K mutants/inNanoluc and HERV-K GagProPol. Five days after transfection, spliced nanoluciferase activity within the retrotransposed HERV-K was measured as the nanoluciferase activity. (B and C) *P* values were determined with Student’s *t* test. \**P* < 0.01; \*\**P* < 0.001; \*\*\**P* < 0.0001; n.s., not significant.

To determine the preferred loci for new integration of HERV-K, we analyzed the new integration sites of HERV-K/inBLC, which encodes an intron-containing blasticidin gene (Fig. 6A), in HeLa cells using a ligation-mediated PCR to amplify the host-virus junction (see Method for more details). We identified total 311 HERV-K LTR integration sites in the genome of HeLa cells (Fig. 6B). Nine of these 311 HERV-K LTR integration sites (1p13.2, K1; 4p16c, K6; 6p21.32; 6q26, K12; 10q24.2b, De12; 11q12.2, K18; 15q13.1; 15q22.2, K24; 19q12, K28) were consistent with previously discovered non-reference HERV-K insertions (Subramanian et al, 2011) and were present in HeLa and other cell lines, including fibroblast cells (data not shown). Six of the non-reference HERV-K LTR integration sites (12p13.31, 16p12.3, 7p22.1, 11q13.4, 11q22.1, and 19p12) were present in HeLa cells but not in other cell lines, such as fibroblast cells (Table 1 Universal in HeLa, Fig. 6B and 6C). One of the six non-reference integration sites was almost identical to one cited in a previous report by John Coffin’s group (19p12b, K113), and the others have not yet been reported. Compared to the universal integration sites in HeLa cells, 21 of the new HERV-K/inBLC integration sites occurred in introns, exons, or intergenic regions (Table 1 Specific in HeLa-inBLC, Fig. 6B, 6D-F and Supplemental Fig. S2A). The clone numbers with some HERV-K integration sites (8q24.22, 16p11.2, and Xp11.23) gradually increased during cell culture (Table 1 Specific, rapid-growth, Fig. 6D), whereas the clone numbers with the other integration sites did not increase (Table 1 Specific, normal and slow-growth, Fig. 6E, and Supplemental Fig. S2A). This suggests that HERV-K integrations at 8q24.22, 16p11.2, and Xp11.23 might promote cell growth. Seven of the integration sites were observed in HeLa cells but not in HERV-K/BLC-transfected HeLa cells (Fig. 6F, Supplemental Fig. S2B). However, all of these sites in HeLa cells were presented in low clone numbers and were detected in regions of repeated sequence, such as short interspersed nuclear elements (SINEs). Since DNA sequence data we obtained in this study is short reads, it is difficult to argue the reliability of integration sites in the repeated sequences. Some of the integrated HERV-K DNAs were amplified by nested PCR with the indicated primers shown in Fig. 6A. The expected amplification products of ∼1000–1500 bp (2F/2R) and ∼3000 bp (5F/5R) were confirmed in HERV-K/inBLC-transfected HeLa cells, but those of ∼1500–2000 bp (3F/2R) were not (Fig. 6G, Supplemental Fig. S2C). It was possible that HERV-K retrotransposition is dependent on the integration machinery through the 3’ poly(A) tail of RNA similar to LINE1 (Doucet et al, 2015). According to our sequencing analysis, HERV-K integrase yielded a 5–6-bp target-site duplication (TSD), which is conserved in the stably integrated provirus, as in alpha-, beta-, gammaretroviruses and lentiviruses but not in LINE1, in the regions flanking the HERV-K integration sites. Moreover, the CMV promoter at the 5’-LTR was replaced with U3 in each integrated HERV-K through reverse transcription (Fig. 6H, Supplemental Fig. S2D). Of note, although transient expression of the transfected HERV-K/inBLC construct driven by the CMV promoter allows one round of retrotransposition, subsequent retrotransposition does not occur in HeLa cells because HERV-K LTR is not activated in the absence of SOX2. In summary, these results indicate that reverse-transcribed HERV-K_CON_ genomes are preferentially integrated into intron and inter-genes through the retroviral integration machinery and potentially influence the cell proliferation depending on the integration sites.

**Table 1.**
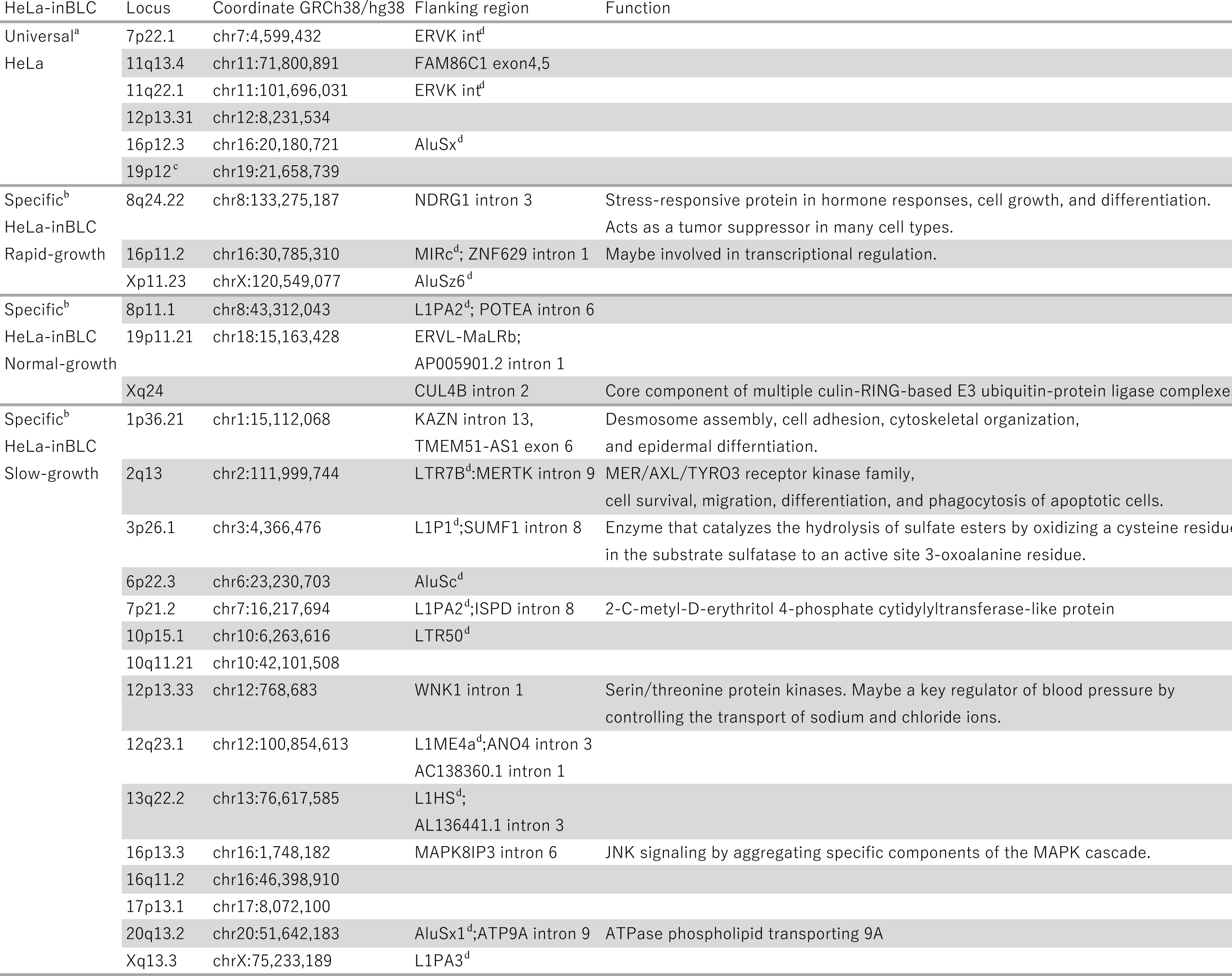
Loci of new HERV-K integration sites in HERV-K-transfected HeLa cells. ^a^Universal HERV-K integration sites in HeLa cells. ^b^Different HERV-K integration sites between HeLa and HERV-K/inBLC-transfected HeLa cells. ^c^Consistent with a previous report from John Coffin’s group (Subramanian et al, 2011). ^d^HERV-K flanking region is in the repetitive sequence.

**Fig. 6.**
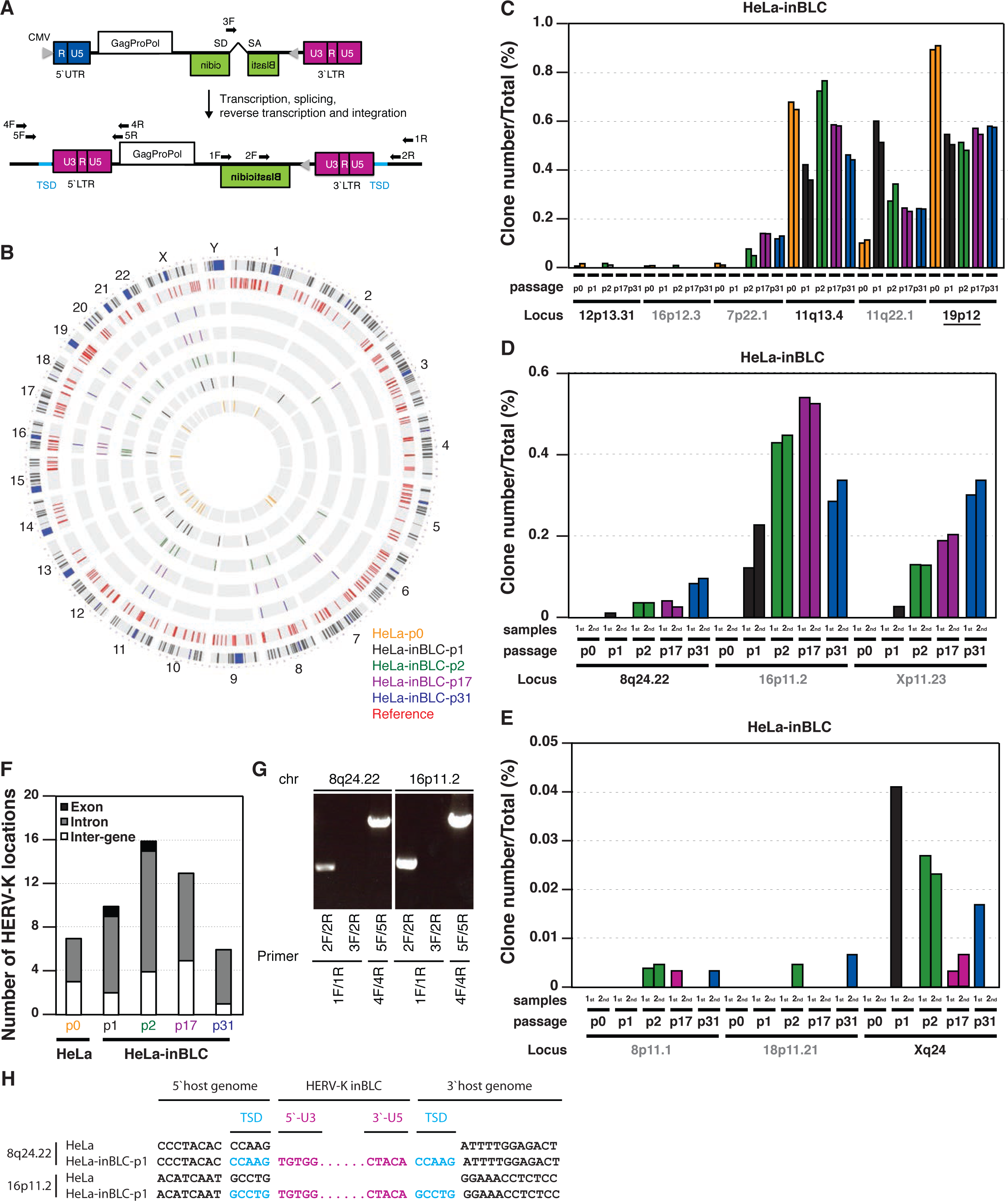
New integration sites for HERV-K appeared in HERV-K-transfected HeLa cells. (A) Construction of pHERV-K GagProPol/inBlasticidin (inBLC) is described. inNluc was replaced with inBlasticidin (inBLC) in the Env region of HERV-K. HeLa cells were transfected with pHERV-K inBLC. Blasticidin-resistant cells were selected 2 weeks after transfection, and the HERV-K DNA in the genome was then amplified by PCR and determined with NGS analysis. The primers were designed to bind the outside of repeated sequences (4F, 5F, 1R and 2R arrowheads), (B) HERV-K integration sites that are present in the database are shown in red (Reference). Non-reference HERV-K integration sites in the HeLa and HeLa-inBLC cells, but not in fibroblasts, are shown in yellow (passage 0), black (passage 1), green (passage 2), purple (passage 17), and blue (passage 31). The outer most ring is G-band of human chromosome. (C) Universal non-reference HERV-K insertions (Universal) were detected in both HeLa and HeLa-inBLC cells, but not in fibroblast cells. (D and E) Specific non-reference HERV-K insertions (Specific) were detected in HeLa-inBLC-p1, -p2, -p17, and -p31 cells, but not in HeLa-p0. Cell growth rates were classified as rapid (D) or normal (E). (C, D and E) Gray letters indicate HERV-K-integration into a LINE-1, SINE, or repeat elements. These data were collected from two independent samples. (F) The number of specific non-reference HERV-K locations in HeLa and HeLa-inBLC-p1, -p2, -p17, and -p31 cells were determined. (G) HERV-K integration sites (8q24.22 and 16p11.2) in HeLa-inBLC-p1 cells were confirmed with PCR. Arrows (A) indicate primer-binding sites for PCR. The primers, which annealed at the integration sites, were designed to sequences adjacent to HERV-K LTR (A). (H) Sequences between HERV-K LTR and the neighboring HERV-K genomes in HERV-K/inBLC-transfected HeLa cells were analyzed with Sanger sequencing. TSD indicates target-site duplication in the human genome generated by integrase.

### Endogenous HERV-K retrotransposition occurs in iPS cells

Recently, iPS cells have become potential research models for regenerative medicine. To develop iPS cells, fibroblast cells are reprogrammed by at least three factors: SOX2, OCT3/4, and KLF4. Therefore, we speculated that HERV-K expression might be induced by SOX2 in iPS cells. As expected, large amounts of HERV-K Gag mRNA were detected in iPS cells compared with NCCIT cells (Supplemental Fig. S3A). It is possible that unregulated HERV-K transposes in the genomes of iPS cells. To investigate HERV-K retrotransposition, we analyzed the HERV-K integration sites in fibroblast cells and iPS cells from the same donor with an NGS analysis (See Method for more details) (Fig. 7A). We found six non-reference HERV-K insertions in both the fibroblast cells and iPS cells, which were not found in HeLa cells (Fig. 7B, Table 2 Universal in fibroblast and iPS cells). Two of the six non-reference HERV-K insertions were consistent with HERV-K integration sites previously reported by John Coffin’s group (12q12, K20; 13q31.3, K22). Other non-reference HERV-K insertion sites might be unique individual-specific HERV-K sites. On the other hand, we detected non-reference HERV-K integration sites in inter-gene, intron and exon in iPS cells but not in fibroblast cells (Table 2 Specific in iPS cells, Fig. 7C and 7D, Supplemental Fig. S3C). There were four non-reference integration sites in the fibroblast cells that were not present in the iPS cells (Table 2 Specific fibroblast and Supplemental Fig. S3B). Compared with the fibroblast cells, the number of non-reference integration sites in the iPS cells increased with time (Table 2 Specific in iPS31, iPS41, Fig. 7C and 7D, Supplemental Fig. S3C). These results indicate that HERV-K has retrotransposition activity and moves within the genomes of iPS cells. However, we found no rapidly growing cells containing non-reference HERV-K integration sites among the iPS-p41 cells (Fig. 7C), indicating that clonal expansion is rare in iPS cells. These results suggest that newly integrated HERV-K is not always advantageous to the cell proliferation, and cellular clonality might depend on the HERV-K integration sites in iPS cells.

**Table 2.** Loci of new HERV-K integration sites in iPS cells. ^a^Universal HERV-K integration sites in this donor. ^b^Different HERV-K integration sites between fibroblast and iPS cells. ^c^Consistent with a previous report from John Coffin’s group (Subramanian et al, 2011). ^d^HERV-K flanking region is in the repetitive sequence.

| iPS | Locus | Coordinate GRCh38/hg38 | Flanking region | Function |
| --- | --- | --- | --- | --- |
| Universal<sup>a</sup> | 3q13.2 | chr3:113,032,475 | HERV-K int<sup>d</sup>; AC078785.1 intron 1 |  |
| fibroblast | 4q25 | chr4:111,308,304 | AluY<sup>d</sup> |  |
| iPS31 | 6q14.1 | chr6:77,725,406 | HERV-K int<sup>d</sup>; MEI4 intron 2 | Required for meiotic DNA double-strand break formation. |
| iPS41 | 7p14.3 | chr7:334,071,494 | L1PA3<sup>d</sup>; BBS9 intron 19 | Required for ciliogenesis but is dispensable for centriolar satellite function. (bardet-biedl syndrome) |
|  | 12q12<sup>c</sup> | chr12:43,919,854 | L1MB1<sup>d</sup>; TMEM117 intron 2 |  |
|  | 13q31.3<sup>c</sup> | chr13:90,090,934 | ; LINC00559 intron 3 | Long non-coding RNA. |
| Specific<sup>b</sup> | 8q13.1 | chr8:66,953,856 | L1PA2<sup>d</sup>; TCF24 intron 3 | Putative transcription factor. |
| fibroblast | 9q22.33 | chr9:99,500,005 | L1PA2<sup>d</sup> |  |
|  | 15q15.1 | chr15:41,053,309 | AluY<sup>d</sup>; INO80 intron 19 | DNA helicase, transcriptional regulation, DNA replication, DNA repair |
|  | 21q11.2 | chr21:13,820,849 | MIR/LTR5Hs<sup>d</sup> |  |
| Specific<sup>b</sup> | 1p13.2 | chr1:113,755,874 | AluY<sup>d</sup>;PHTF1 intron 2 | Transcription regulation |
| iPS31 | 2q22.3 | chr2:147,700,152 |  |  |
|  | 5p13.3 | chr5:30,822,734 | L1PA2<sup>d</sup> |  |
|  | 15q11.2 | chr15:23,715,336 | L1PA3<sup>d</sup> |  |
|  | 20q11.23 | chr20:38,348,370 | AluSc5<sup>d</sup>; LBP intron 1 | Binds to the lipid A moiety of bacterial lipopolysaccharides (LPS), a glycolipid present in the outer membrane of all Gram-negative bacteria. (toxic pneumonitis, mastitis, infective endocarditis, alcoholic hepatitis, mesenteric lymphadenitis, peritonitis, appendicitis, bacteremia, endocarditis, acute respiratory distress syndrome) |
|  | Xq25 | chrX:126,189,98 | L1PA2<sup>d</sup> |  |
| Specific<sup>b</sup> | 1p13.2 | chr1:111,609,710 | AluSx<sup>d</sup>; RAP1A intron 1 | Induces morphological reversion of a cell line transformed by a Ras oncogene. (Kabuki syndrome 1, leukocyte adhesion deficiency type iii, tuberous sclerosis, babesiosis, central nervous system hemangioma, cerebral cavernous malformations-1, cerebral cavernous malformations-2) |
| iPS31 |  |  |  |  |
| iPS41 |  |  |  |  |
|  | 1p34.3 | chr1:38,842,086 | AluY<sup>d</sup>; RRAGC intron 6 | Has guanine nucleotide-binding activity |
|  | 1p35.1 | chr1:32,614,846 | AluSc<sup>d</sup> |  |
|  | 3q11.2 | chr3:97,642,693 | L1PA2<sup>d</sup>; EPHA6 intron 14 | (Oculoauricular syndrome) |
|  | 4q22.1 | chr4:91,979,478 | L1HS<sup>d</sup> |  |
|  | 5p13.3 | chr5:32,212,988 | AluSx3<sup>d</sup> |  |
|  | 9q21.13 | chr9:73,987,206 | /LTR5Hs<sup>d</sup> |  |
|  | 10q24.2 | chr10:99,256,369 | MSTD int<sup>d</sup> |  |
|  | 11q13.2 | chr11:67,868,810 | LTR5Hs int<sup>d</sup> |  |
|  | 11q13.3 | chr11:70,471,356 | ; SHANK2 exon 2 | Molecular scaffolds in the postsynaptic density of excitatory synapses. (Autism susceptibility 17, autism spectrum disorder, pervasive developmental disorder, deafness, autosomal recessive 63, secretory diarrhea, autistic disorder) |
|  | 15q14 | chr15:36,348,199 |  |  |
|  | 16p12.3 | chr16:18,135,887 | AluSz<sup>d</sup>; AC010601.1 intron 1 |  |
|  | 21p11.2 | chr21:8,586,021 | MLT1A0<sup>d</sup> |  |
|  | 21q22.3 | chr21:43,711,450 | AluY<sup>d</sup> |  |

**Fig. 7.**
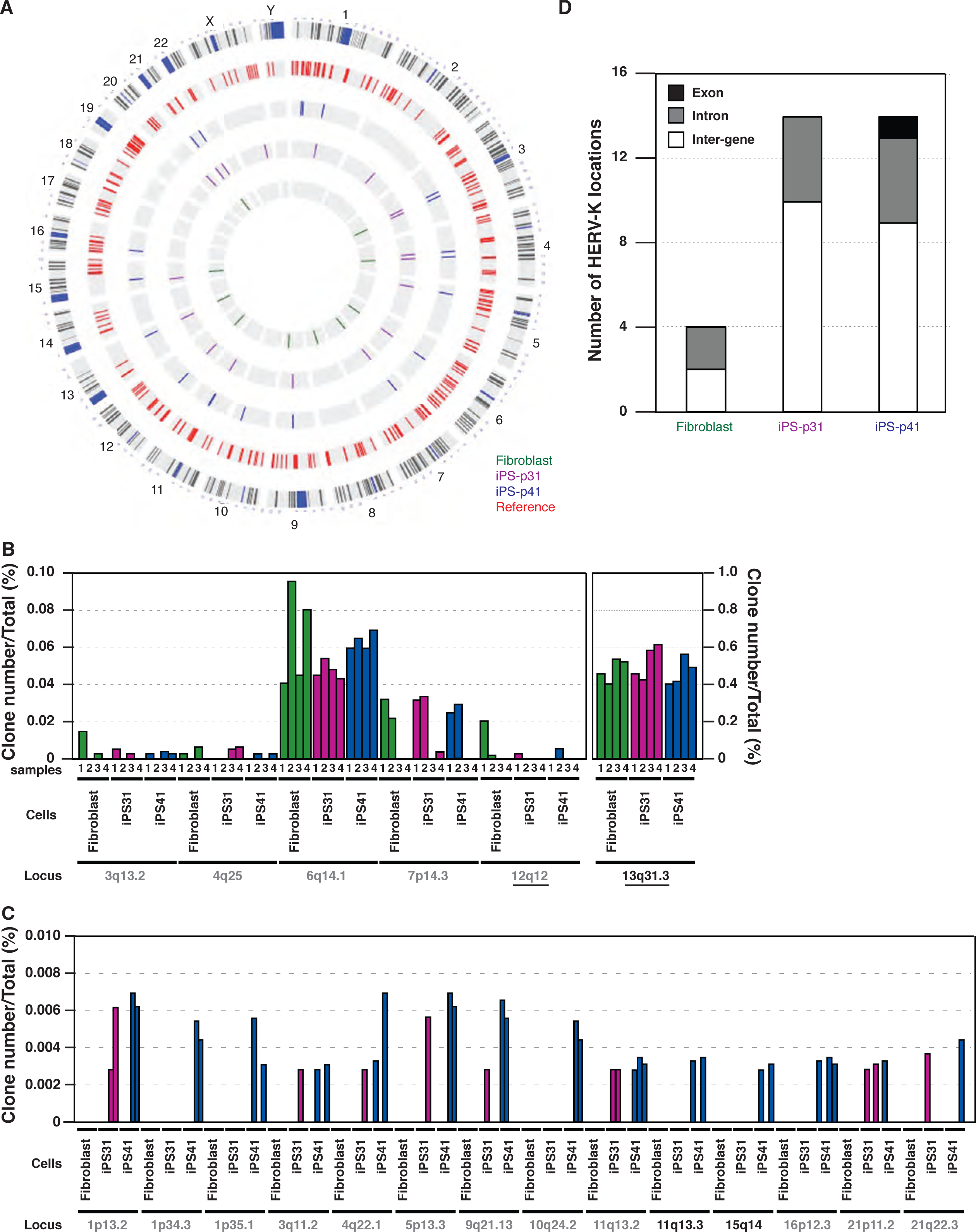
New integration sites for HERV-K appeared in iPS cells. (A) Integration sites of HERV-K were determined with NGS analysis. iPS cells were derived from fibroblasts, then the cells were passaged 31 times (iPS-p31) or 41 times (iPS-p41). HERV-K integration sites that were already present in the database (Reference) are shown in red. Non-reference HERV-K integration sites in fibroblasts and iPS cells but not in HeLa cells are shown in green (fibroblast), purple (iPS cells; passage 31), and blue (iPS cells; passage 41). (B) Universal non-reference HERV-K insertions were specifically determined in this donor. (C) Specific non-reference HERV-K insertions were specifically identified in iPS cells. (B and C) Gray letters indicate HERV-K-integration into LINE-1, SINE, or repeat elements. These data were collected from four independent samples. (D) The number of specific non-reference HERV-K locations in fibroblasts and iPS cells were determined.

## Discussion

In this study, we demonstrated that HERV-K is capable of retrotransposition in SOX2-expressing cells. The transactivation of HERV-K LTR5Hs and LTR5B by SOX2 is retained even after the accumulation of several mutations in these LTR sequences. Although the physiological roles of HERV-K are still unknown, we found that HERV-K has retrotransposition activity and moves randomly around the host genome. In a blasticidin-selected cell population where SOX2 is not expressed, and hence retrotransposition occurs only once after transfection, the copy number of HERV-K, which is integrated into the intron of a tumor suppressor gene (*NDRG1*) (Table 1), increased. This suggests that HERV-K integration may accelerate cell growth by impairing the host genome and thus can cause several diseases (discussed below). However, in SOX2-expressing cells, only a small number of novel HERV-K integration sites were identified. It is possible that the SOX2-expressing cells that have new integration of HERV-K may die or grow slowly during the long-term culture due to the harmful impact of the HERV-K integration on the genome integrity.

In addition to the well known role in the maintenance and re-establishment of pluripotency (Avilion et al, 2003) (Takahashi & Yamanaka, 2006), SOX2 is essential for central nervous system (CNS) development and the maintenance of neural stem cells (Pevny & Nicolis, 2010). SOX2 is also expressed in Schwann cells (Le et al, 2005) and impairs Schwann cell remyelination and their functional recovery after nerve injury, such as in multiple sclerosis (Roberts et al, 2017). Therefore, it is conceivable that SOX2-induced expression of HERV-K might have impact on CNS development, the maintenance of neural stem cells, remyelination, or recovery from nerve injury. Indeed, HERV-K is implicated in several neural diseases, including multiple sclerosis (Tai et al, 2008). Moreover, the HERV-K LTR integration sites differ slightly among the genomes of individual humans and between human tissues, and HERV-K LTR single-nucleotide polymorphisms (SNPs) are implicated in several neural diseases (Wallace et al, 2018). It is possible that SOX2 might influence the expression of genes adjacent to HERV-K LTR5Hs. Additionally, our results are consistent with the possibility that HERV-K expression, which becomes uncontrollable when the epigenetic regulation of SOX2 is disturbed, disrupts the nervous system through the retrotransposition of HERV-K.

SOX2 is known for its association with numerous types of cancer (Weina & Utikal, 2014). It regulates the self-renewal and maintenance of cancer stem cell populations by promoting oncogenic signaling (Bareiss et al, 2013; Chen et al, 2012; Laga et al, 2011). The expression of HERV-K is considerably higher in malignant tissues, such as germ-cell tumors, melanomas, and ovarian cancers, than in healthy tissues (Buscher et al, 2005; Conrad et al, 1997; Kurth & Bannert, 2010; Wang-Johanning et al, 2007), suggesting a possibility that the transactivation of HERV-K LTR5Hs by SOX2 is involved in numerous malignant tumors. Whether HERV-K expression is involved in the self-renewal and maintenance of cancer stem cells is still unknown, but it is possible that the impairment of the genome by HERV-K retrotransposition may cause the malignancy of tumor tissues.

HERV-K is transiently reactivated in early human development to protect cells from the threat of exogenous viral infection (Grow et al, 2015); however, HERV-K retrotransposition entails a risk of genomic impairment in SOX2-expressing cells such as iPS cells. According to our results, such genomic impairment is probably a rare event in iPS cells (Fig. 7). In addition to the possibility that HERV-K retrotransposition causes a defect in cell growth, thereby reducing cells with the genomic impairment, it is possible that HERV-K retrotransposition is prohibited by host restriction factors during its reverse transcription and/or integration. For example, APOBEC3F, a restriction factor in cell-free HERV-K infection (Lee & Bieniasz, 2007), may inhibit HERV-K retrotransposition during the reverse transcription step. In the yeast *Saccharomyces cerevisiae*, Ty1 LTR retrotransposon Gag forms virus-like particle as retrosome for the reverse transcription (Salinero et al, 2018). APOBEC3G interacts with Ty1 Gag in the retrosome and restricts the Ty1 retrotransposition (Dutko et al, 2005; Schumacher et al, 2005). However, it is unknown whether HERV-K Gag forms a retrosome, as does Ty1 (Salinero et al, 2018), or whether APOBEC3F can access the HERV-K genome in the retrosome. In future, the mechanism of HERV-K retrotransposition must be clarified.

Transposable elements such as HERVs often provide new functions to vertebrate hosts, which result in the exaptation (Johnson, 2019). Endogenous retrovirus Env, which is called syncytins, is necessary for fusion of cytotrophoblasts to form the multinucleate syncytiotrophoblast layer of the placenta (Lavialle et al, 2013). The syncytins are involved in the convergent evolution during the changes from oviparity to viviparity because syncytins are originated independently across multiple mammalian lineages and a live-bearing reptile (Cornelis et al, 2017; Cornelis et al, 2015). Additionally, the neuronal gene Arc, which is retrotransposon Gag protein, mediates intercellular signaling in neurons and is essential for the animal cognition (Pastuzyn et al, 2018). It suggests that retrotransposon Gag has obtained alternative functions in neurons during evolution. The function of HERV-K is still unclear, but considering the acquisition of SOX2 responsive elements long time ago and the retention of the competent elements in their LTRs since then, it is tempting to speculate that HERV-K play important physiological roles in SOX2-expressing cells.

## Acknowledgments

We would like to thank Dr. T. Kadomatsu and the members of Dr. Satou’s laboratory for their technical advice on the ChIP assay and the Illumina MiSeq sequencing analysis. We would also like to thank Dr. Paul D. Bieniasz for providing plasmids. This work was supported by MEXT KAKENHI (Grants-in-Aid for Scientific Research under grant number 15K21242 and grant number 20K07517) to K.M.; the Takeda Science Foundation to K.M. We thank Janine Miller, PhD, from Edanz Group (www.edanzediting.com/ac), for editing a draft of this manuscript.

## Author’s Contributions

KM, YM, and TS conceived and coordinated the study. KM, SY, TM and HT performed the experiments. YS and YU supported the NGS analysis. MG, TK, KT, and TE prepared the iPS cells. FW supported the preparation of the knockout cells.

JI and II prepared the phylogenetic trees. AO supported the writing of the manuscript. All authors read and approved the final manuscript.

## Declaration of Interests

The authors declare that they have no competing interests.

### Plasmids

Full-length HERV-K_CON_ was kindly provided by Paul Bieniasz (Lee & Bieniasz, 2007). pHERV-K_CON_ LTR-Luc encodes the luciferase gene, driven by HERV-K LTR. pMXs-SOX2, OCT3/4, KLF4, and NANOG were obtained from Addgene. CHKCinNluc and CHKCinBLC were derived from CHKCP (kindly provided from Paul Bieniasz) (Lee & Bieniasz, 2007). The puromycin N-acetyl-transferase gene was removed from CHKCP (CHKCP/delPuro), and a *Not*I site was inserted. Intron-disrupted Nanoluc (inNluc) and blasticidin (inBLC) were designed as previously reported (Xie et al, 2011). The inNluc and inBLC cassettes encode the SV40 early enhancer/promoter and SV40 late poly(A) signal, respectively. These cassettes were introduced into CHKCP/delPuro at the *NotI* site in an antisense orientation.

### Cells

HeLa cells were cultured in Dulbecco’s modified Eagle’s medium (DMEM; Sigma) supplemented with 5% fetal bovine serum (FBS). NCCIT cells (ATCC^®^ CRL-2073™) were cultured in RPMI1640 medium with 10% FBS, 1 mM sodium pyruvate and Glutamax™ (Teshima et al, 1988). Human iPS cells were generated from human-skin-derived fibroblasts and were cultured with mitomycin-C-treated mouse embryonic fibroblast feeder cells, as described previously (Fusaki et al, 2009; Soga et al, 2015); Eto et al., 2018).

### HERV-K retrotransposition assay

HeLa cells were seeded in six-well plates at a density of 2 x 10^5^ cells/well. The cells were transfected with Lipofectamine 3000 Reagent (Invitrogen), according to the manufacturer’s protocol. The cells were harvested 1– 6 days after transfection, and the nanoluciferase activities in the cells were measured with Nano-Glo® Luciferase Assay Reagent (Promega).

### Measurement of dual-luciferase luminescence

Luminescence was measured with the Dual-Luciferase Reporter Assay System (Promega), according to the manufacturer’s instructions. The cell lysate was mixed with Luciferase Assay Reagent II. Firefly luciferase activity was measured with a luminometer. *Renilla* luciferase activity was read after the cell lysate containing Luciferase Assay Reagent II was mixed with Stop & Glo Reagent.

### Bisulfite sequencing

EpiTect Plus Bisulfite conversion kit (Qiagen) was used as described previously (Grow et al, 2015). The PCR fragments were inserted into pCR-BluntII-Topo vector (Invitrogen). Approximately 10 clones in HeLa and NCCIT cells were Sanger sequenced for quantifying the CpG methylation.

### ChIP assay

NCCIT cells were fixed with 1% formaldehyde and lysed with 20% NP-40 (10 mM HEPES-KOH pH 7.9, 1.5 mM MgCl_2_, 10 mM KCl, 0.5 mM DTT, and 20% NP-40 with protease inhibitor cocktail [Roche]). The chromatin in the lysates was fragmented to 320 bp after digestion with micrococcal nuclease. After further lysis with 10% SDS (50 mM Tris-HCl pH 8.1, 0.2 mM EDTA, 10% SDS, with protease inhibitor cocktail [Roche]), the chromatin was sonicated. The DNA–protein complexes were precipitated overnight by incubation with an anti-SOX2 antibody (BioLegend),and then incubated with ChIP-Grade Protein G Magnetic Beads (9006, Cell Signaling Technology) for 2 h. The abundance of HERV-K LTR in the precipitated-DNA was analyzed with quantitative PCR using primers: HERV-K LTR primer F 5’-AGCACTGAGATGTTTATGTG-3’ and R 5’-TGTGGGGAGAGGGTCAGC -3’ and SYBR Premix ExTaq II (Takara Bio Inc.). The signal intensity was quantified with the ABI 7900HT Fast Real-Time PCR System (Applied Biosystems).

### Linker-mediated (LM)-PCR

The HERV-K integration sites were analyzed with LM-PCR and high-throughput sequencing, as previously described (Gillet et al, 2011; Satou et al, 2017). To analyze HERV-K integration site, the junction between the 3’-LTR of HERV-K and the host genomic DNA was amplified with a primer targeting the 3’-LTR and the linker. The first forward primer targeting the 3’-LTR was B3-K1: 5’-CCTCCATATGCTGAACGCTGGTT-3’; the second forward primer targeting the 3’-LTR was P5B5-K2: 5’-AATGATACGGCGACCACCGAGATCTACACCCAAATCTCTCGTCCCACCTTACGA GAAACACCCACAGG-3’. The DNA libraries were sequenced as paired-end reads with Illumina MiSeq, and the resulting fastq files were analyzed. The sequencing primer targeting the 3’-LTR was Seq-K1: 5’-ACACCCACAGGTGTGTAGGGGCAACCCACC-3’. The flanking host genome sequences were used to determine viral integration sites. The resulting short reads were cleaned using an in house script, which extracts reads with a high index-read sequencing quality (Phred score > 20) in each position of an 8-bp index read. The clean sequencing reads were aligned with HERV-K LTR sequences (CCTACA and CCTTCA) and were mapped to human genome using the BWA-MEM algorithm (Li & Durbin, 2009). Further data processing and cleanup, including the removal of reads with multiple alignments and duplicated reads, were performed using Samtools (Li & Durbin, 2009) and Picard (http://broadinstitute.github.io/picard/). The clone numbers with each HERV-K integration site were quantified as previous reports (Gillet et al, 2011; Satou et al, 2017).

### Western blotting analysis

Cells and virus were lysed with 1% Triton X lysis buffer (50 mM Tris-HCl pH 7.5 containing 0.5% Triton X-100, 300 mM NaCl, 10 mM iodoacetamide, and protease inhibitor cocktail [Roche]). After 2 x SDS sample buffer was added, the SOX2, OCT3/4, GAPDH and HERV-K Gag proteins were detected with immunoblotting using an anti-SOX2 antibody (Merck Millipore), an anti-OCT3/4 antibody (BD Biosciences), an anti-GAPDH antibody (Sigma) and an anti-HERV-K Gag antibody (Austral Biologicals), respectively, as the primary antibodies. Horseradish peroxidase (HRP)-conjugated anti-mouse Ig antibody (Jackson ImmunoResearch) was used as the secondary antibody. The HRP-conjugated secondary antibody was detected with Chemi-Lumi One L (Nacalai Tesque).

### Reverse transcription–quantitative PCR analysis

Total RNA was purified with the RNeasy Mini Kit (Qiagen). The mRNA was reverse transcribed with *Murine leukemia virus* (MLV) reverse transcriptase after it was annealed with a poly(T) primer. HERV-K *gag* DNA was amplified with primers: HERV-K Gag CA forward primer 5’-CAAGACCCAGGAAGTACCT-3’ and reverse primer 5’-ACACTCAGGATTGGCGTT-3’. All qPCR assays were performed with SYBR Premix ExTaq II (Takara Bio Inc.). The data for the target genes were then normalized to the expression level of glyceraldehyde 3-phosphate dehydrogenase (*GAPDH*; housekeeping gene), amplified with GAPDH forward primer 5’-CGCTCTCTGCTCCTCCTGTT-3’ and reverse primer 5’-ACAAAGTGGTCGTTGAGGGC-3’.

### Flow cytometry analysis

Total DNA was extracted from NCCIT cells with the DNeasy Blood & Tissue kit (Qiagen). HERV-K LTR was amplified with HERV-K LTR primers (5Hs forward primer: 5’-CCAAAAGCCATCGATTGTGGGGAAAAGCAAGAGAG-3’; 5Hs 5’LTR reverse primer: 5’-TTCCATCTCGAGTGAAGTGGGGCCAGCCCCTCCACACCT-3’; 5Hs 3’LTR reverse primer: 5’-TTCCATCTCGAGTGTAGGGGTGGGTTGCCCCTCCACACC-3’; 5A forward primer: 5’-AAAGCCATCGATTGTAGGGAAAAGAAAGAGAGATCAGAC-3; 5A 5’LTR reverse primer: 5’-TTCCATCTCGAGTGAAGGGGTGGCCTGCCCCTCCA-3’; 5A 3’LTR: reverse primer 5’-TTCCATCTCGAGCTCCACACCTGTGGGTAT-3’; 5B forward primer: ‘-AAAGCCATCGATTGTAGGGAAAAGAAAGAGAGATCAG-3’; 5B 5’LTR reverse primer: 5’-TTCCATCTCGAGTGAAGTGGGGCCAGCCCCTCCACACCT-3’; 5B 3’LTR reverse primer: 5’-TTCCATCTCGAGCTCCACACCTGTGGGTATTTCT-3’). The HERV-K LTR was inserted upstream from the yellow-fluorescent-protein-encoding gene (HERV-K LTR-Venus). HeLa and NCCIT cells were cotransfected with pHERV-K LTR-Venus and pMXs-SOX2. Two days after transfection, the fluorescent signals were analyzed with flow cytometry.

## Figure legends

**Fig. S1**. SOX2 and OCT3/4 contribute to the promoter function of HERV-K LTR. (A) SOX2- and OCT3/4-binding motifs were identified in the HERV-K LTR with the PROMO software, which is used to identify putative transcription factors. Deletion mutants of LTR were constructed. (B–D) NCCIT cells were transfected with the pHERV-K_CON_ mutants. Two days after transfection, the firefly luciferase activities were measured with a luciferase reporter assay. (E) HeLa cells were cotransfected with pHERV-K_CON_ mutants and the indicated plasmids. Two days after transfection, the firefly luciferase activities were measured with a luciferase reporter assay. (F) HeLa cells were cotransfected with the pHERV-K_CON_ mutants, the indicated plasmids, and the Renilla-Luc plasmid. Two days after transfection, both firefly luciferase and *Renilla* luciferase activities were measured with a dual luciferase reporter assay.

**Fig. S2**. New integration sites for HERV-K appeared in HERV-K-transfected HeLa cells. (A and B) HeLa cells were transfected with pHERV-K-inBLC. Blasticidin-resistant cells were selected 2 weeks after transfection, and the HERV-K DNA in the genome was then amplified with PCR and analyzed with next-generation sequencing. Specific non-reference HERV-K insertions (Specific) were detected in HeLa (B) or HeLa-inBLC-p1, -p2, -p17, and -p31 cells (A). Cell growth speeds was classified as slow (A). Gray letters indicate HERV-K-integration in LINE-1, SINE, or repeat elements. (C) HERV-K integration sites (1q36.21 and 12p13.33) in HeLa-inBLC-p1 cells were confirmed with PCR. (H) Sequences between HERV-K LTR and the neighboring HERV-K genomes in HERV-K/inBLC-transfected HeLa cells were analyzed with Sanger sequencing. TSD is a target-site duplication in the human genome generated by integrase..

**Fig. S3**. New integration sites of HERV-K appeared in iPS cells. (A) HERV-K Gag mRNA and GAPDH mRNA expression in NCCIT and iPS cells was measured with reverse transcription–PCR. The integration sites of HERV-K were determined with next-generation sequencing. (B and C) Specific non-reference HERV-K insertions were specifically determined in fibroblasts (B) and iPS-31 cells (C). Gray letters indicate HERV-K-integration in LINE-1, SINE, or repeat elements.

